# Dexamethasone Enhances CAR T Cell Persistence and Function by Upregulating Interleukin 7 Receptor

**DOI:** 10.1101/2022.08.29.505735

**Authors:** Ashlie Munoz, Ryan Urak, Ellie Taus, Claire Hsieh, Katherine Jin, Shu-Hong Lin, Dennis Awuah, Vibhuti Vyas, Saul J. Priceman, Mary C. Clark, Stephen J. Forman, Xiuli Wang

## Abstract

Dexamethasone (dex) is a glucocorticoid that is a mainstay for treatment of inflammatory pathologies, including immunotherapy-associated toxicities. Dex suppresses the endogenous immune response and is also believed to suppress the function of chimeric antigen receptor (CAR) T cells. However, recent reports observed higher CAR T cell numbers in patients treated with dex, highlighting the rationale for interrogating the specific effects of dex on CAR T cells. Here, we found that dex did not inhibit CAR T cell expansion or function. A single dose of dex during the manufacturing process upregulated the pro-persistence interleukin 7 receptor α (IL7Rα) on CAR T cells and induced expression of genes involved in activation, migration, and persistence. The *ex vivo* upregulation of IL7Rα induced by dex significantly enhanced CAR T cell persistence and anti-tumor efficacy *in vivo* when combined with exogenous IL-7. Moreover, the combination of dex and IL-7 resulted in increased persistence of CAR T cells and led to complete remission of mice. Overall, our studies in both *in vitro* and *in vivo* treatment support a positive role of dex on CAR T cell potency and provide insight into the application of glucocorticoids in cellular anti-cancer therapy.

## Introduction

Chimeric antigen receptor (CAR) T cells have revolutionized the treatment of hematological malignancies and have yielded a staggering 70-90% response rate against acute lymphoblastic leukemia (ALL) (1-4). Response and durability of response to CAR T cell therapy is affected by CAR T cell function and persistence following infusion, which are in turn influenced by variables related to attributes of the CAR construct and manufacturing process used to generate the CAR T cells (5). Manufacturing platforms of CAR T cells fundamentally influence *in vivo* longevity and persistence by altering the differentiation state-fitness of T cells (5, 6); more differentiated T cells have shorter persistence relative to memory T cells. Thus, CAR T cell products can be optimized to be enriched for a memory phenotype that is associated with superior potential for anti-tumor efficacy and persistence by altering the manufacturing process.

One method to enrich for a T cell memory phenotype is by leveraging the interleukin 7 (IL-7) pathway. IL-7 is a cytokine that binds to IL-7Rα (CD127) and initiates signaling through the JAK-STAT pathway, which plays a critical role in T cell homeostatic proliferation and memory formation (7). IL-7Rα is highly expressed on naïve and central memory T cells, but found in low levels on effector memory T cells (7, 8). In CAR T cell therapy, patients who have increased levels of serum IL-7 following lymphodepleting chemotherapy have increased progression free survival and overall survival, suggesting that IL-7 signaling may enhance the durability of CAR T cell therapy (9, 10). However, CAR T cells lose expression of IL-7Rα due to activation and *ex vivo* expansion during the manufacturing process (11-13), which may subsequently impair *in vivo* persistence (13). To overcome this problem, our lab and others previously investigated constitutively active IL-7 receptor constructs that confer exogenous, cytokine independent, cell-intrinsic STAT5 signaling and resulted in improved adoptive T cell therapies in pre-clinical models (14-16). However, these methodologies are not readily translatable to the clinic to due toxicity concerns. Thus, establishing clinically applicable methods of retaining and/or upregulating IL-7Rα expression either during manufacturing or post-CAR T cell infusion may result in CAR T cells that are enriched for memory phenotype and capable of enhanced anti-tumor efficacy and persistence.

Glucocorticoids (GCs) are a class of steroids that are routinely used to treat inflammatory conditions in the context of cancer, autoimmunity, and COVID-19 (17). Binding of GCs to the glucocorticoid receptor (GR) causes GR dimerization and translocation to the nucleus, where GRs can modulate expression of genes that contain a glucocorticoid response element (GRE) (5-9), including those involved in T cell proliferation and survival. In general, GCs are not routinely given to patients receiving CAR T cells, except to treat severe neurotoxicity associated with CAR T cell therapy, due to the concern that GCs may dampen CAR T cell activity and persistence (18, 19). However, recent evaluation of clinical data suggests that treatment with GCs, including the synthetic GC dexamethasone (dex), does not affect proliferation or persistence of CAR T cells (20). The effect of GC treatment during CAR T cell therapy has not been directly examined and the role GCs may play on CAR T cell function and persistence have not been described. Intriguingly, there is a GRE upstream of IL7Rα promoter and GC treatment has been shown to upregulate IL-7Rα expression on both murine and human T cells (21, 22). Therefore, we hypothesized that GC treatment may enhance CAR T cell function by upregulating the expression of IL-7Rα.

With the goal of understanding the effect of GCs on CAR T cells, especially potential positive regulation, we evaluated the effects of dex on CAR T cells *in vitro* and *in vivo*. We have found that dex upregulated IL-7Rα on T cells during all stages of T cell differentiation. CAR T cells manufactured in the presence of dex upregulated IL-7Rα and had an upregulation in gene pathways associated with activation, cytotoxicity, and memory. CAR T cells treated with dex had superior anti-tumor efficacy and persistence compared to untreated CAR T cells *in vivo* in a xenograft mouse model of ALL. Moreover, tumor-bearing mice treated with dex and IL-7 post-CAR T cell infusion had superior survival outcomes compared to control mice. Overall, this work identifies a novel role for GCs in potentiating CAR T cell function and persistence through the upregulation of IL-7Rα.

## Results

### Dex does not affect CAR T cell proliferation or effector function

To investigate how dex may affect CAR T cells, we treated CAR T cells with various concentrations of dex (0.1, 1, or 10μM), either once or 3 times (every 48 hours) during the cell expansion phase of *ex vivo* manufacturing and assessed CAR T cell expansion and function (**Figure 1A**). CAR T cell expansion was not impacted by a single dose of dex at any of the concentrations examined (**Figure 1B**). While CAR T cell expansion was not affected by 3 doses of either 0.1 or 1µM dex, 3 doses of 10µM dex decreased CAR T cell expansion relative to control CAR T cells (**Figure 1C**). We therefore adopted 1μM dex for all subsequent experiments. To rule out cell population effects, we examined the impact of 1µM dex on the expansion of CAR T cells generated using either peripheral blood mononuclear cells (PBMC), a widely used T cell population for generating clinical CAR T cells, or naïve/memory T cells (Tn/mem), our clinical platform at City of Hope. Dex did not impact expansion of either PBMC or Tn/mem-derived CAR T cells (**Figure S1A-B**), even when given 3 times during *ex vivo* expansion when compared to untreated CAR T cells (**Figure S1C**). Moreover, dex did not impact the growth of CAR T cells with different target specificities (CD19 or CS1) (**Figure S1B)**, suggesting that dex does not affect the expansion of CAR T cells in general.

**Figure 1.**
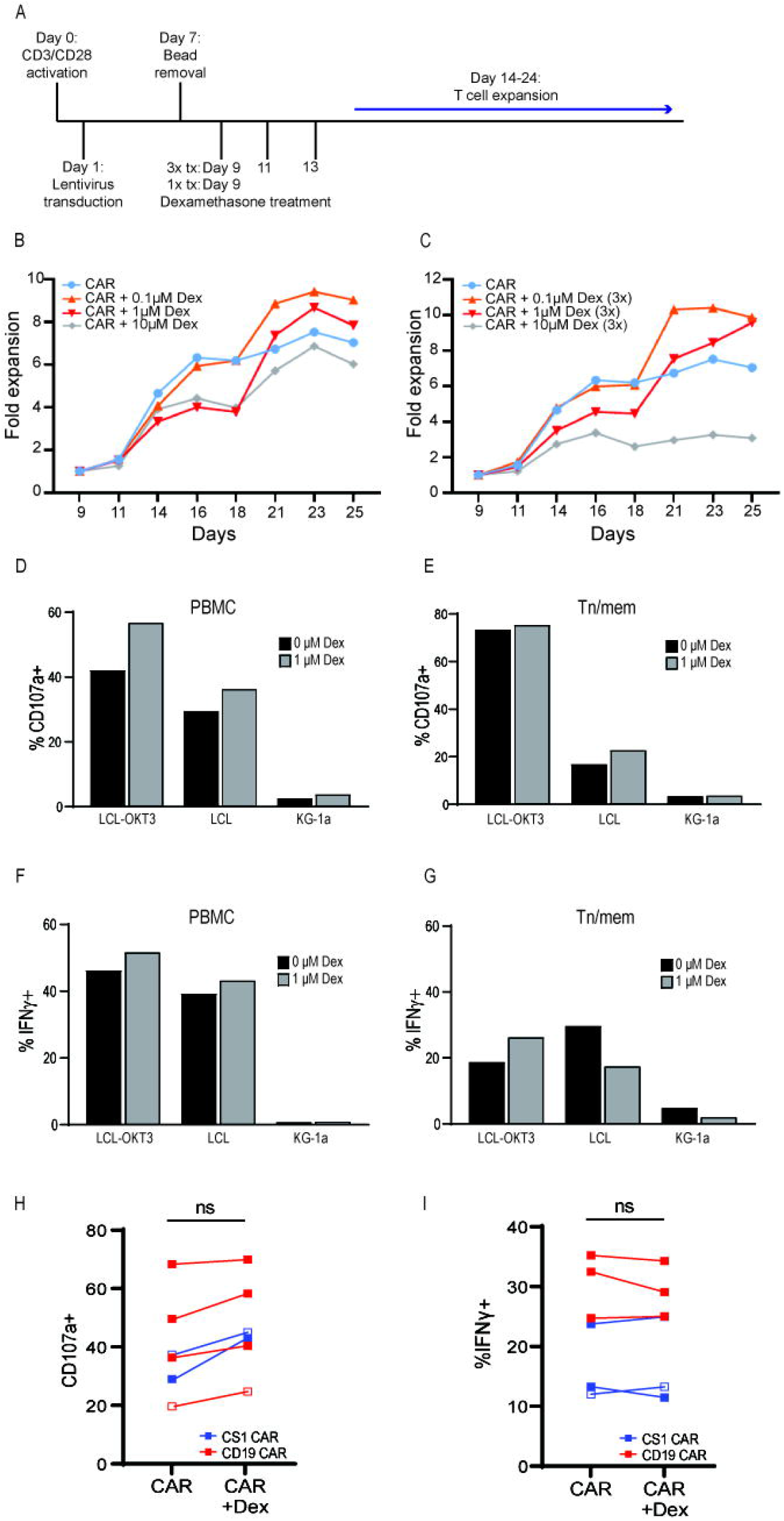
Dex does not affect CAR T cell proliferation or effector functions. **(A)** PBMCs or purified Tn/mem cells were activated with anti-CD3/anti-CD28 beads on day 0, transduced with lentivirus on day 1, followed by bead removal on day 7. CAR T cells were either treated with a single dose of dex on day 9 or three doses on days 9, 11, and 13. CAR T cells were then expanded through days 14-24. Proliferation of Tn/mem-derived CAR T cells from the same donor grown in the presence or absence of a single **(B)** or three **(C)** doses of various concentrations (0.1, 1, 10µM) of dex are depicted. Representative data of CD107a degranulation by PBMC-derived (n=4) **(D)** or Tn/mem-derived (n=2) **(E)** CD19-CAR T cells after 6 hours co-culturing with CD19-positive target LCL cells are presented. Intracellular IFNγ secretion by PBMC-derived (n=5) (**F**) or Tn/mem-derived (n=1) (**G**) CD19 CAR after overnight stimulation with LCL cells. Myeloid leukemia KG-1a was used as a negative control and OKT3-expressing LCL was used as a positive control. **(H)** Accumulative data of CD107a degranulation by PBMC (n=4, closed symbols) or Tn/mem (n=2, open symbols) CS1 (blue) or CD19 (red) CAR T cells after 6 hours of co-culturing with target cells. **(I)** Accumulative data of intracellular IFNγ secretion by PBMC (n=5, closed symbols) or Tn/mem (n=1, open symbols) CS1 (blue) or CD19 (red) CAR T cells after overnight stimulation with target cells.

To determine whether dex impacts CAR T cell effector function, we evaluated dex-treated CD19-CAR T cells for the ability to degranulate and produce cytokines upon co-culture with CD19+ target LCL cells. When compared to untreated CAR T cells, dex-treated CAR T cells had comparable degranulation (**Figure 1D-E, H**) and cytokine production (**Figure 1F-G, I**) as determined by CD107a and INFγ positivity, respectively. This phenomenon was also not CAR-specific, as the effector functions of both CD19- and CS1-CAR T cells were unaffected by dex (**Figure 1H-I**). To determine if this observation was dose dependent, we co-cultured CAR T cells with 0.1, 1, or 10μM of dex and found that even at high concentrations there was no impact on CAR T cell function (**Figure S1D**). Together, these data suggest that CAR T cells are not impacted by dex exposure during the cell expansion phase of manufacturing.

### Dex upregulates IL7Rα on CAR T cells

We hypothesized that dex could affect the expression of IL7Rα on CAR T cells, as the IL7Rα gene contains a GRE and dex increases IL7Rα expression on both mouse and human T cells (21, 22). Therefore, we monitored IL7Rα expression on CAR T cells treated with or without dex over the course of culture: before activation (D0), immediately after transduction and bead removal (D9), and 7 days after culture with or without dex (D16), for both PBMC-(**Figure 2A**) and Tn/mem-derived (**Figure 2B**) CAR T cells. Consistent with our previous studies (12), ∼80% of pre-activated PBMC-derived T cells were positive for IL7Rα, while only ∼50% of T cells were IL7Rα positive following activation/transduction (**Figure 2A**). However, following dex treatment, ∼80% of CAR T cells expressed IL7Rα, while IL7Rα expression decreased to ∼40% in untreated CAR T cells (**Figure 2A**). This trend was consistent in Tn/mem-derived CAR T cells (**Figure 2B**) and was independent of CAR construct (**Figure S2A-B**) or dex dosage (**Figure S2C**). Together, these data suggest that dex mediates upregulation of IL7Rα on CAR T cells.

**Figure 2.**
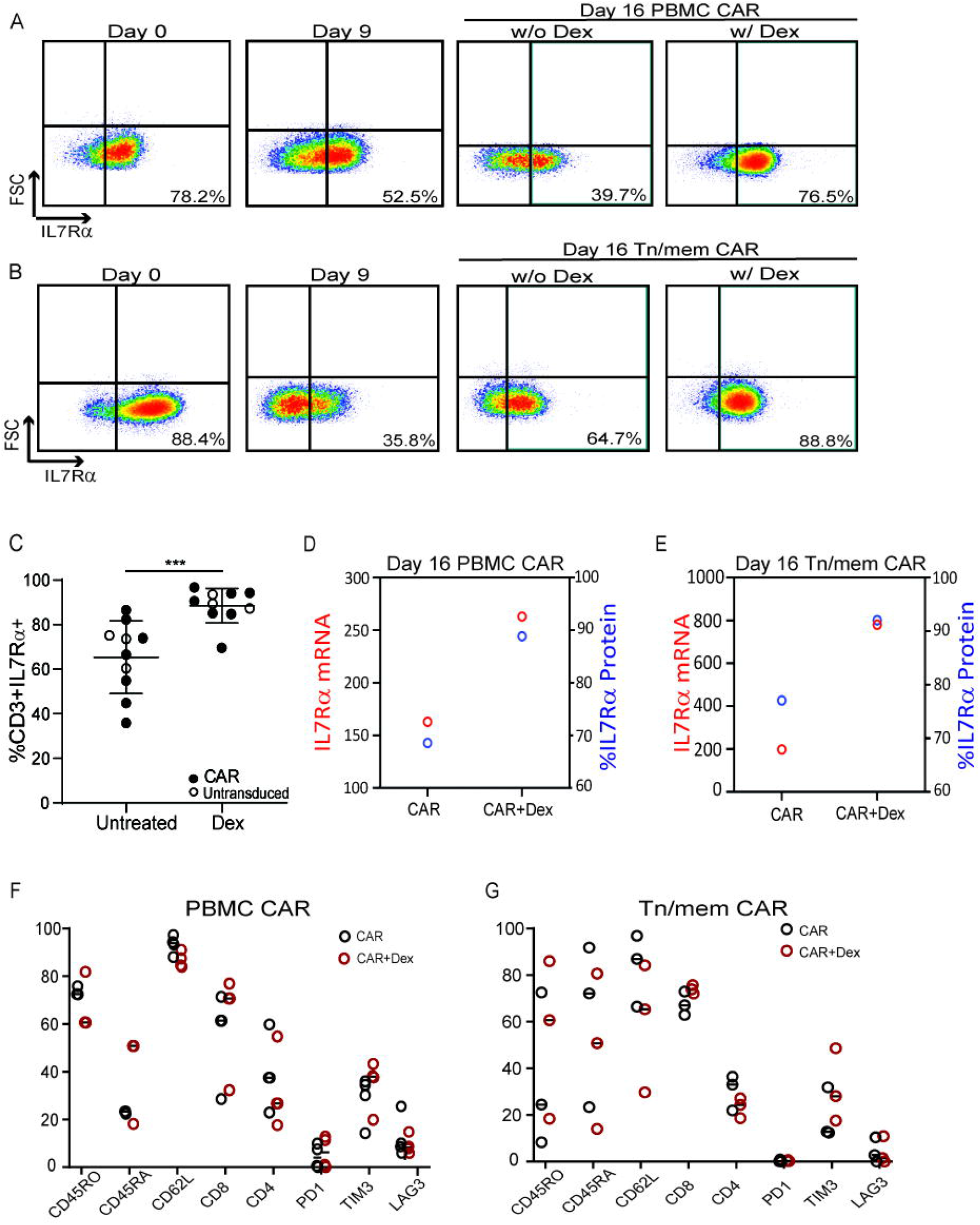
Dex upregulates IL7Rα expression by CAR T cells. IL7Rα surface expression on PBMC-derived **(A)** and Tn/mem-derived **(B)** CD19-CAR T cells prior to transduction (Day 0), after transduction (Day 9) and after dex treatment (Day 16). Data are either representative of seven separate donors **(A)**, four separate donors **(B)** or cumulative data (p=0.00005) **(C)**. IL7Rα mRNA and protein expression in PBMC-derived **(D)** and Tn/mem-derived **(E)** CAR T cells, treated with or without 1μM dex. Phenotypic characterization of PBMC-derived (n=12) **(F)** and Tn/mem-derived (n=6) **(G)** CAR T cells in the treated with or without 1μM dex is presented.

To understand dex-mediated IL7Rα upregulation on T cells, we assessed the ability of dex to increase IL7Rα on extensively expanded and differentiated, Epstein Barr virus (EBV)-specific central memory (Tcm) and effector memory (Tem) T cells, the latter of which being a population with low to no expression of IL7Rα (7, 8). We expanded the EBV-specific Tcm and Tem cells *ex vivo* for 3 months by rapid expansion method (REM), treated with or without dex and analyzed surface IL7Rα (23). Both Tcm and Tem virus-specific T cells had increased IL7Rα expression upon dex treatment (**Figure S2D**), suggesting that dex can mediate IL7Rα expression on T cells regardless of differentiation status. Additionally, we determined that IL7Rα was upregulated at both protein and mRNA levels following treatment with dex (**Figure 2C-E**), providing evidence that dex regulates IL7Rα expression at the transcriptional level in T cells. To investigate whether dex induced other changes that affect CAR T cell function, we subjected CAR T cells treated with or without dex to phenotypic analysis by flow cytometry. We observed similar levels of memory markers and exhaustion features on dex treated vs. untreated CAR T cells **(Figure 2F-G and Figure S2A-B)**, which was consistent regardless of CAR construct **(Figure S2A-B)**. Furthermore, we analyzed IL2 alpha, beta, and gamma chain receptor expression after dex treatment and found IL7Rα was the only gamma chain cytokine receptor upregulated by dex (**Figure 3**). Together, these data suggest that dex specifically modulates IL7Rα expression on CAR T cells.

**Figure 3.**
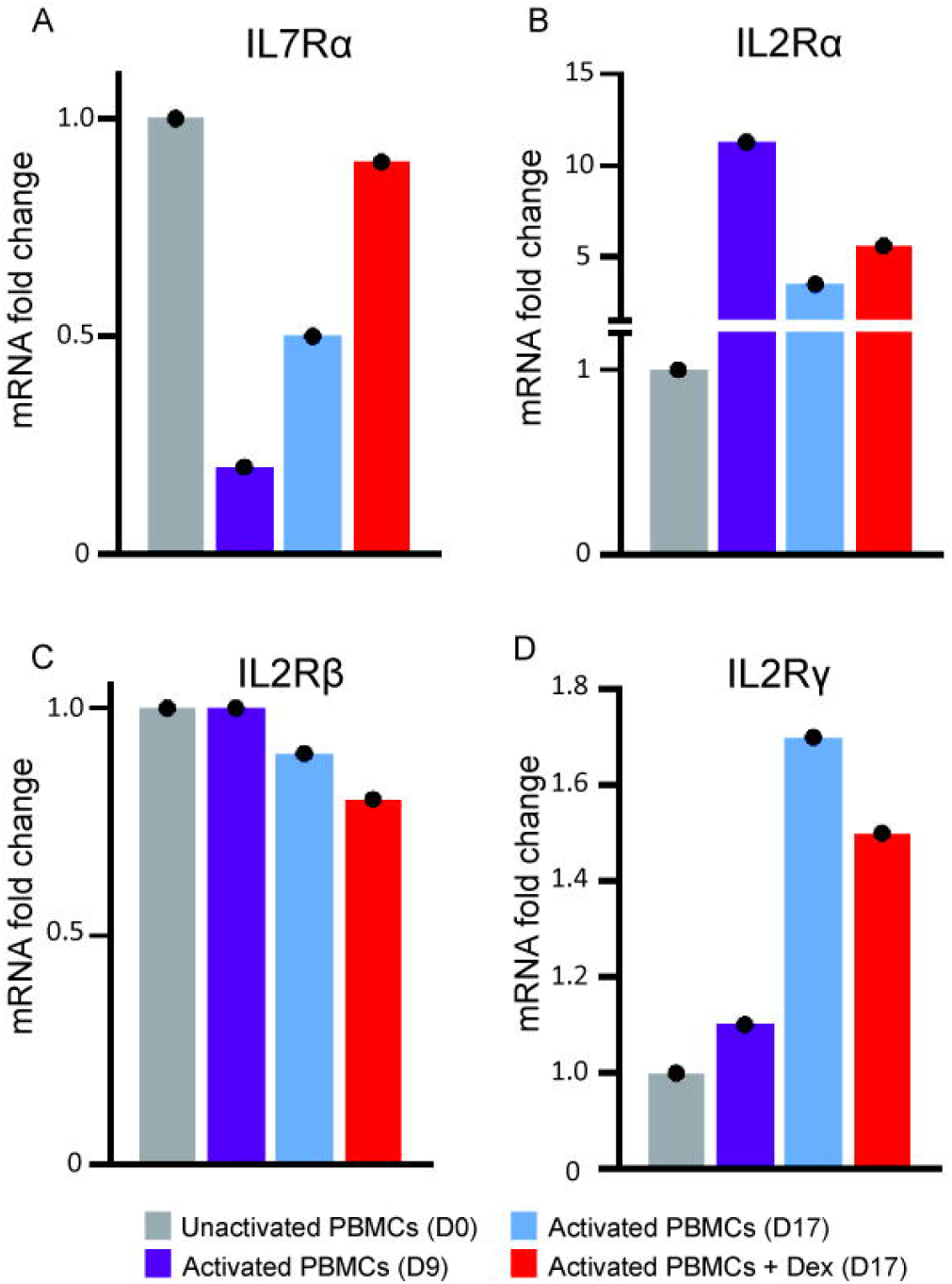
Dex treatment does not influence other gamma chain cytokine receptors. Levels of mRNA expression of IL7Rα **(A)**, IL2Rα **(B)**, IL2Rβ **(C)**, and IL2Rγ **(D)** on resting (Day 0), activated (Day 9), and expanded (Day 17; 7 days in dex) PBMC-derived CD19-CAR T cells, treated with and without dex, were analyzed by NanoString gene analysis.

We next interrogated how dex affected CAR T cells at the transcript level by performing NanoString analysis. Briefly, PBMC and Tn/mem-derived CAR T cells were expanded for 16 days (7 days in dex) and analyzed for changes in gene signatures and signaling pathways (**Figure 4**). In pathway score analysis, we observed upregulation in pathways related to activation, migration, persistence, and chemokine production in dex-treated CD19-CAR T cells relative to untreated CD19-CAR T cells. In contrast, pathways related to apoptosis and TCR diversity were downregulated after dex treatment (dashed lines indicate z > +1.96 or z < −1.96) (**Figure 4A-D**). We identified 33 genes that were differentially regulated by dex in both PBMC- and Tn/mem-derived CAR T cells, including IL7Rα, relative to housekeeping genes. Dex upregulated activation-related genes such as IFNGR2, VAV3 (24), DDIT4 (25) and IL-31 (26), migration gene CXCR4 (27), and AREG (28), as well as genes that promote memory T cell formation, such as TNFRSF11A and ACVR1C (29) (**Figure 4E-F**). To determine whether the changes in gene expression induced by dex were long-lasting, we analyzed PBMC- and Tn/mem-derived CAR T cells treated with or without dex that were expanded for 23-24 days after initiation of CAR T cell manufacturing (i.e., ∼14 days after dex treatment). We observed that following extended *in vitro* culture, many pathways and genes that were previously observed to be up-or down-regulated (**Figure 4)** returned to a comparable state as untreated CAR T cells for both populations (PBMC and Tn/mem-derived CAR T cells), suggesting that the effects of dex were reversible (**Figure 4, Figure S3**).

**Figure 4.**
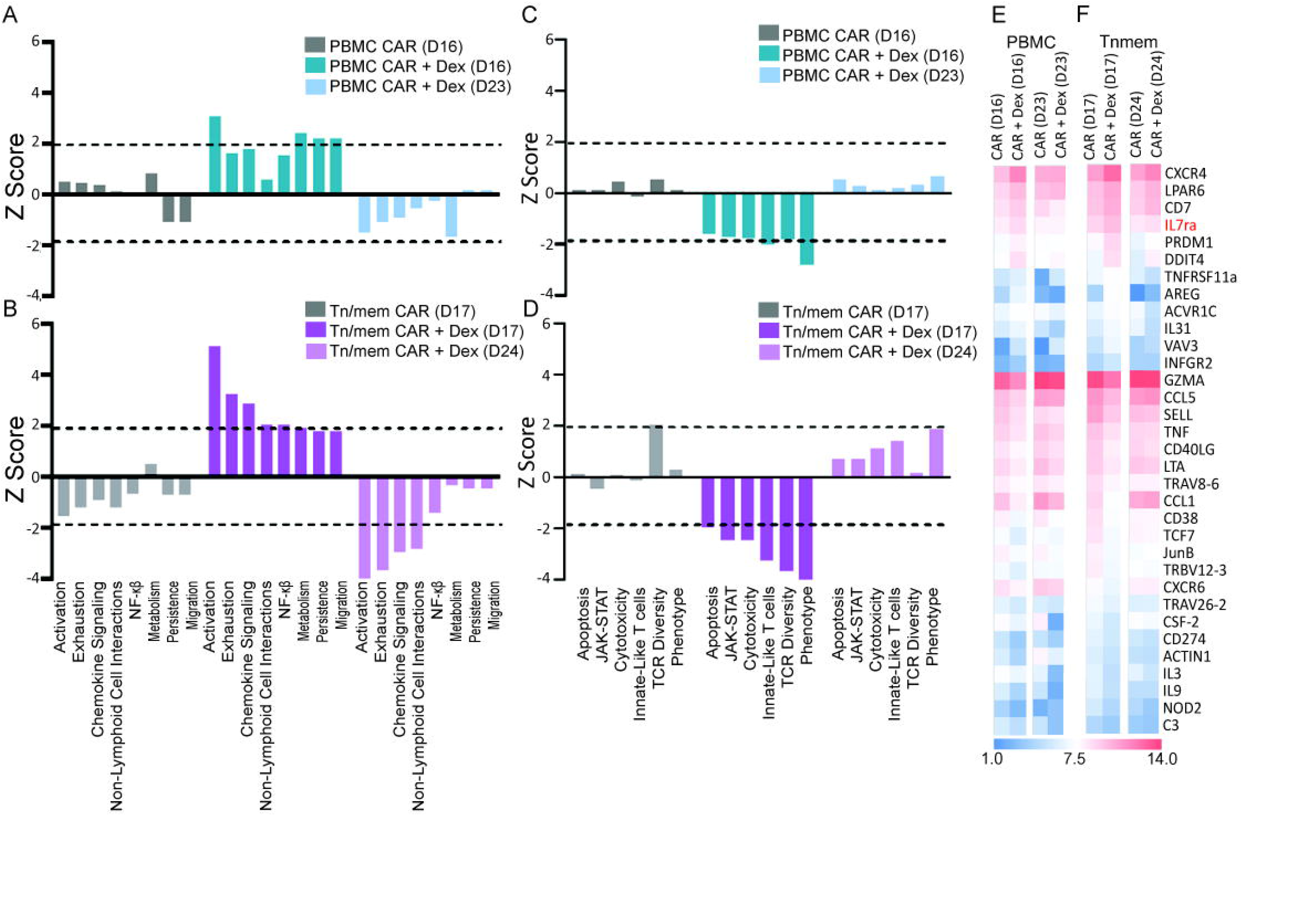
Dex regulates signaling and gene expression of CAR T cells. CAR T cells treated with and without dex were collected on Day 16 and 23 for PBMC-derived CAR T cells and Day 17 and 24 for Tn/mem-derived CAR T cells. Pathway (**A-D**) and gene expression (**E-F**) analysis were performed with NanoString. nCounter Advanced Analysis software provided by the manufacturer were applied for pathway score analysis. The dash lines indicated z = +/-1.96 **(A-D)**. For gene expression trend analysis, genes were first grouped into upregulated genes, downregulated genes and genes with no response based on the changes of normalized gene expression counts between cells treated with and without dex. Only genes within the same groups in both PBMC-derived and Tn/mem-derived CAR T cells were included. Genes with minor changes were excluded based on the %CV cutoff value of housekeeping genes. The genes shown on the heatmap is on a log scale with base 2 **(E-F)**.

### Dex treated CAR T cells are more responsive to IL7

To determine whether dex-induced IL7Rα expression on CAR T cells conferred biological activity, we analyzed the expansion of CAR T cells treated with various concentrations of dex 1 or 3 times as previously described and supplemented with IL7 (**Figure 5**). CAR T cells grown in the presence of IL7 had higher expansion when compared to CAR T cells not given IL7, regardless of dex dosing. Interestingly, CAR T cells that were treated with 3 doses of 10μM dex, a condition that we previously observed to have decreased expansion compared to untreated cells (**Figure 1C**), had greater expansion in the presence of IL7 than without IL7 (**Figure 5B**). This data suggests that potentially the negative effect that dex had on CAR T cell expansion at high concentrations was overcome when the IL7 pathway is triggered.

**Figure 5.**
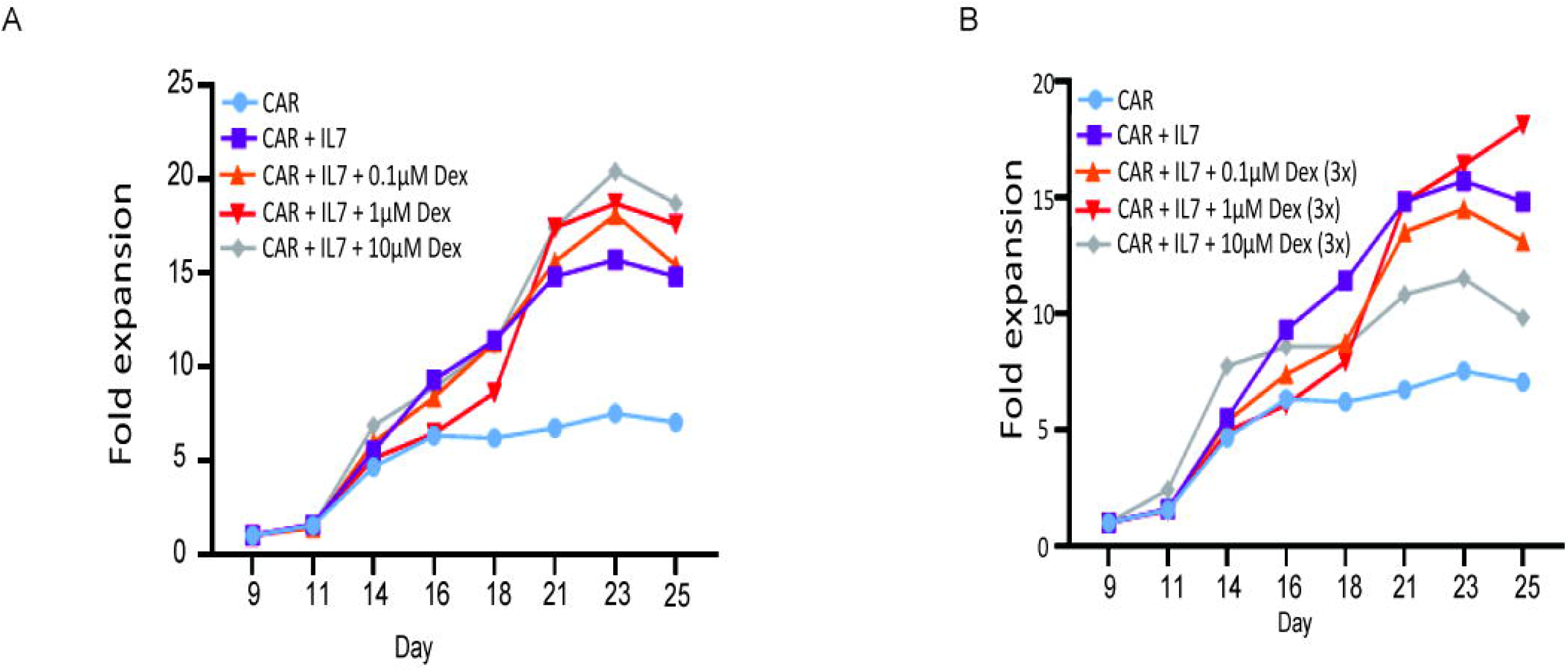
IL7 enhances expansion of dex-treated CAR T cells. Proliferation of Tn/mem CAR T cells grown in the presence or absence of a single **(A)** or three **(B)** doses of various concentrations (0.1, 1, 10µM) and 10ng/mL IL7, supplemented every other day. CAR T cells without dex and IL7 were used as controls.

To test the antitumor activity of dex-treated CD19-CAR T cells *in vivo* (**Figure 6A**), we engrafted 0.5×10^6^ ALL cells (SupB15) into NOD-ScidIL2Rγnull (NSG) mice and allowed 5 days for engraftment. We treated tumor bearing mice with 1×10^6^ PBMC-derived CD19-CAR T cells that were generated with or without 1μM dex *ex vivo* (**Figure 6B**) and monitored tumor burden and survival. To simulate the increased serum IL7 observed clinically in a subset of patients following lymphodepletion (9, 10), we injected 10×10^6^ irradiated, human (hu)IL-7-secreting CHO cells intraperitoneally (i.p.) every other day for a total of 6 doses, starting 24 hours post CAR T cell injection. We confirmed an increase in huIL-7 in the serum of huIL-7 CHO treated mice (**Figure S4**). The combination of CAR T cells manufactured with dex and huIL-7 resulted in significantly better antitumor activity, as seen at Day 55, than either CAR T cells manufactured in dex alone or CAR T cells alone (P<0.01), as well as the untreated group (P<0.001) (**Figure 6C-D**). Mice treated with CAR T cells manufactured with dex and huIL-7 also had significantly prolonged survival (P<0.001) when compared to untreated mice and mice treated with both CAR T cells and huIL-7 injections (**Figure 6E**). This data supports the *in vivo* biological activity of IL7Rα upregulated by dex. To confirm the data is not CAR and/or tumor dependent, we conducted the same experiment with CS1-CAR T cells in multiple myeloma model and consistent data were observed (**Figure S5)**. In line with the reversible mechanism of dex *in vitro* (**Figure 4**), when we treated SupB15-engrafted mice with CAR T cells that had been expanded for 23 days, we observed inferior potency (**Figure S6**) vs. CAR T cells that were expanded 16 days *ex vivo* (**Figure 6**).

**Figure 6.**
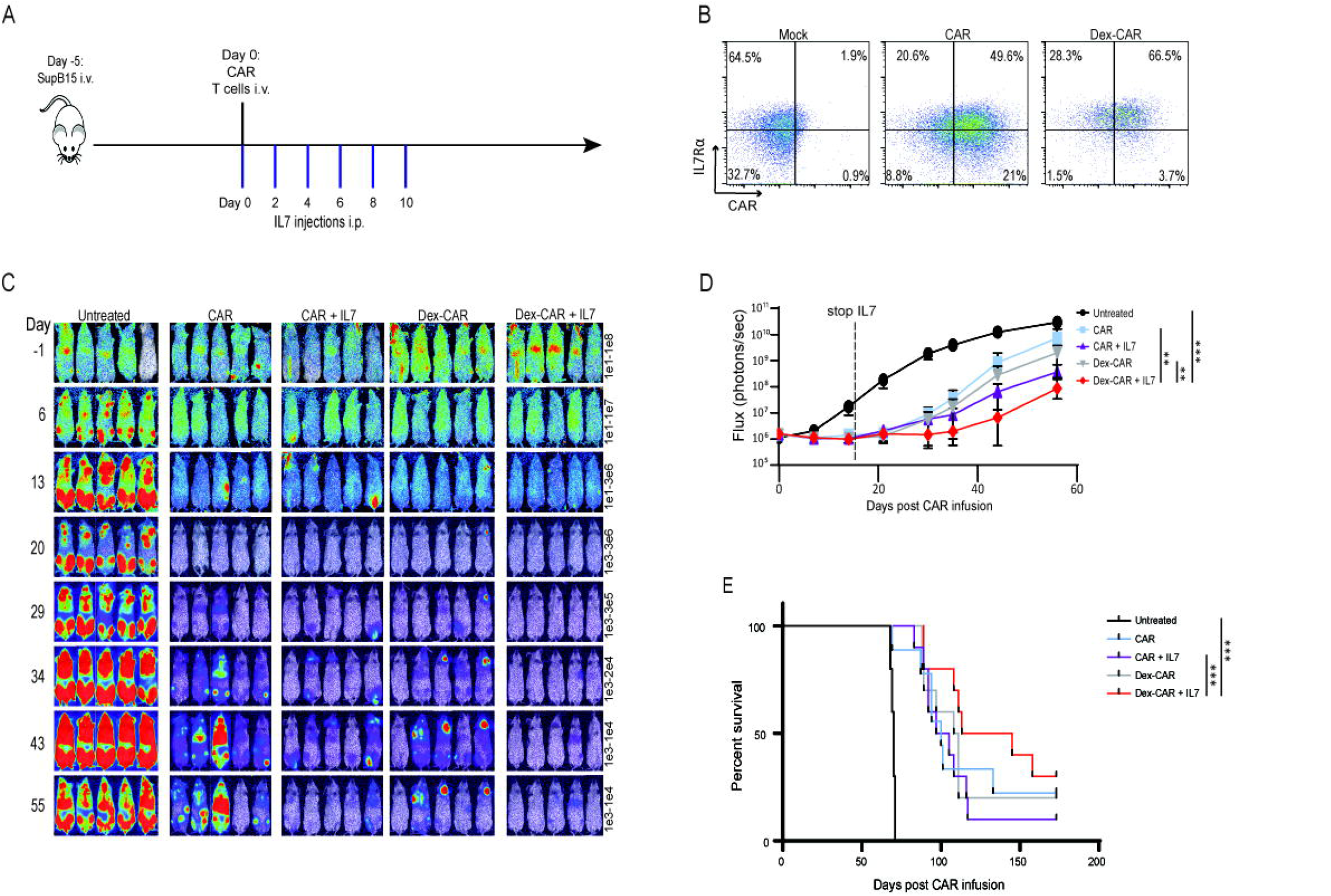
Dex-treated CD19-CAR T cells exhibit improved antitumor activity in a leukemia mouse model. 0.5 × 10^6^ acute lymphoblastic leukemia cell line (SupB15) cells expressing GFP and firefly luciferase (GFPffluc+) were inoculated into NSG mice intravenously (i.v.) on Day −5. After confirmation of engraftment, 1 × 10^6^ CAR T cells harvested 7 days after 1x 1µM Dex treatment were adoptively transferred into tumor-bearing mice i.v. Mice were injected intraperitoneally (i.p.) with 8000 rads-irradiated human IL-7 producing CHO cells (10 × 10^6^) every 48 hours for six injections **(A)**. CAR and IL7Rα expression on input cells **(B)**. Biophotonic imaging was used for tumor signal monitoring **(C)** and tumor burden was measured in Flux (photons/sec) by bioluminescent imaging and was evaluated weekly **(D)**. Kaplan-Meier survival curve **(E)**. N=10 mice per group and images of five representative mice are presented. **P<0.01, ***P<0.001.

### Dex in combination with IL-7 enables CAR T cells to eliminate tumor *in vivo*

Because dex did not impact expansion or effector function of CAR T cells when given during the cell expansion phase of *ex vivo* manufacture, we next interrogated how direct injections of dex and huIL-7 affected the efficacy of untreated CD19-CAR T cells *in vivo* (**Figure 7**). We inoculated NSG mice i.v. with 0.5×10^6^ SUP-B15 cells and treated with a single infusion of 1×10^6^ central memory T cell-derived CD19-CAR T cells (**Figure 7B**) 5 days post-tumor engraftment. We dosed the mice with 10mg/kg dex and 10×10^6^ irradiated huIL-7-expressing CHO cells every other day for the first month and then weekly for 4 additional months. While CD19-CAR T cells and dex alone had some anti-tumor activity compared to untreated mice, the combination of CD19-CAR T cells with dex and huIL-7 completely eliminated the tumors (**Figure 7C-D**) and led to significantly prolonged survival (**Figure 7E**).

**Figure 7.**
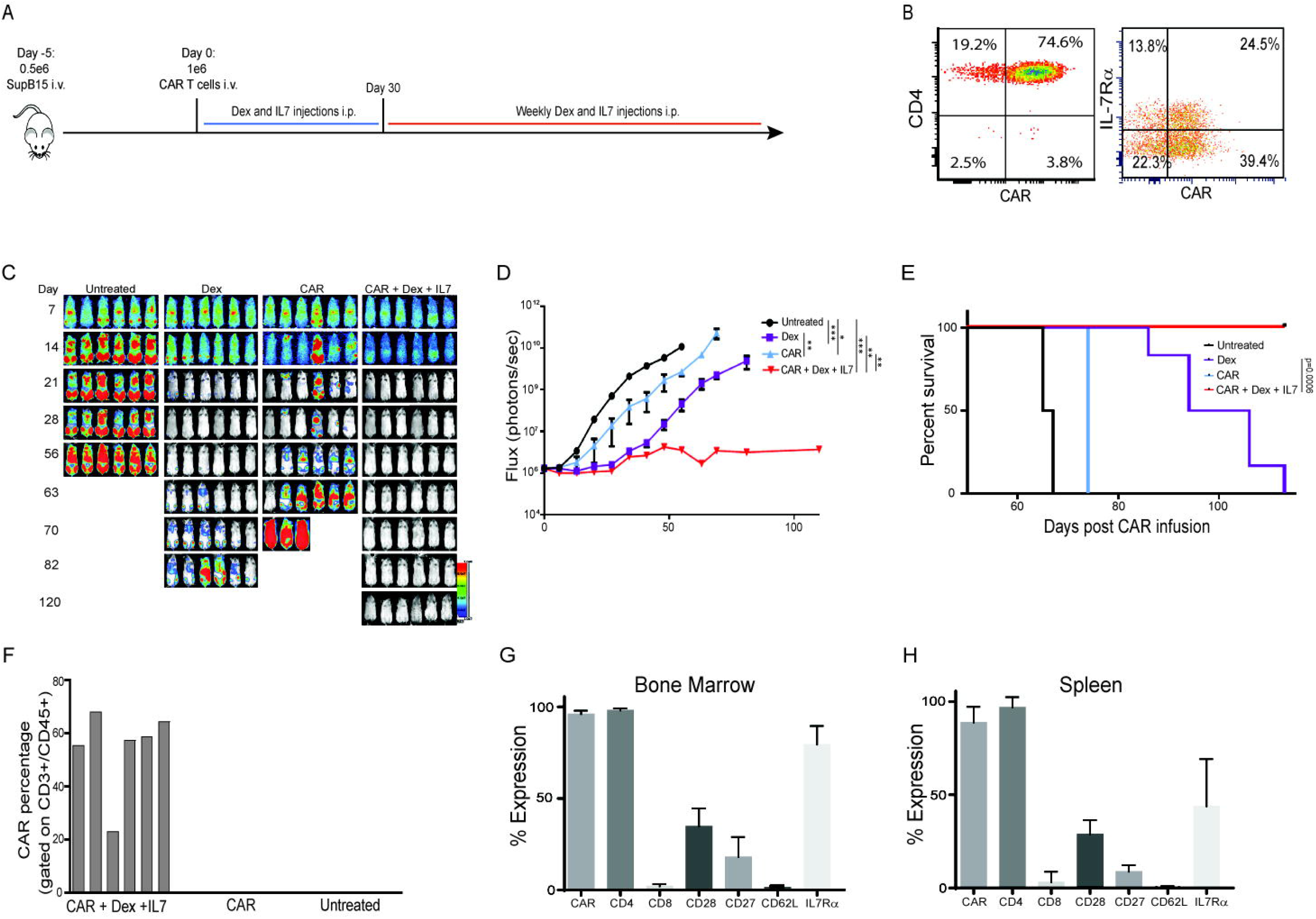
Dex in combination with IL-7 enables CAR T cells to eliminate tumor *in vivo*. Mice (n=6) were engrafted with 0.5 × 10^6^ SUP-B15GFPffluc+ cells i.v. on Day −5. 1 × 10^6^ CD19-CAR T cells were injected i.v. Dex was given 10mg/kg s.c. every other day for 4 weeks and then once a week afterwards. 10 × 10^6^ 8000 rads-irradiated human IL7 secreting CHO cells were injected i.p. every other day for 4 weeks and then once a week afterwards **(A)**. CAR expression on input cells **(B)** Biophotonic tumor imaging for tumor signal and tumor burden was measured in Flux (photons/sec) by bioluminescent imaging and was evaluated weekly **(C, D)**. Kaplan-Meier survival curve **(E**). On Day 21, blood was collected through retroorbital bleeding and analyzed for CAR expression by flow cytometry **(F)**. CAR and immune receptors on gated CD45+CD3+ T cells in lymph organs such as bone marrow **(G)** and spleen **(H)** were harvested on euthanasia analyzed with flow cytometry. **P<0.01, ***P<0.001.

To study CAR T cell persistence after the combination of dex and IL7, we collected blood from mice on Day 21 and assessed for the presence of CAR T cells by flow cytometry. Only mice that received dex, huIL7 and CAR T cells had circulating CAR T cells at this time point (**Figure 7F**), suggesting that IL7/dex increased the persistence of CAR T cells. When we euthanized the mice treated with CAR T cells in combination with dex/huIL7, which occurred 150 days post CAR T cell treatment, we observed high levels of CAR T cells that expressed IL7Rα in bone marrow and spleen (**Figure 7G-H)**. The CAR alone group did not have persistent CAR T cells in either the bone marrow or spleen at euthanasia. Interestingly, persisting CAR T cells had the same proportion of CAR expression as input cells as well as high levels of IL7Rα even after 150 days *in vivo* without tumor. To further confirm these findings, in a separate experiment, we tested the combination therapy with different controls. Consistently, CAR T cells in combination with dex and IL7 was the only group that had no tumor progression (p<0.01) (**Figure S7)** and superior CAR T cell persistence.

## Discussion

Dex is a well-established immunosuppressive agent that is widely believed to suppress the function of CAR T cells. However, recent clinical observations have shown that patients treated with dex following CAR T cell therapy did not have decreased CAR T cell numbers when compared to patients who did not receive dex (20, 28). We evaluated the effect of dex on CAR T cells during the cell expansion phase of *ex vivo* manufacture and found that a single dose of dex did not impact CAR T cell expansion (**Figure 1B-C, Figure S1A-C**) or function (**Figure 1D-G, Figure S1D**), even at a dose (10µM) that was higher than the high dose used clinically to treat patients with CAR T cell associated toxicity (20mg=50nM) (20, 30). These results are in line with previous preclinical studies that show activated T cells are not inhibited by dex (10, 21, 30, 31), which could be attributed to CAR T cell *ex vivo* manufacturing platform leading to IL-2 induced resistance to glucocorticoids (32). We demonstrated that dex upregulated the expression of pro-proliferation and pro-survival gene, IL7Rα (**Figure 2**) at both the mRNA and protein levels (**Figure 4**), regardless of CAR design (**Figure S2A-C**) or T cell differentiation state (**Figure S2D**). When we analyzed gene pathways affected by dex, we found that pathways associated with promoting memory formation, activation, and trafficking were upregulated in CAR T cells following treatment with dex (**Figure 4)**. Together, these findings support a potential beneficial role for dex during CAR T cell therapy, involving multiple positive modulatory pathways, primarily the IL7R pathway.

Because dex increased the expression of IL7Rα on CAR T cells, we analyzed whether dex during the cell expansion phase could yield CAR T cells with superior *in vivo* anti-tumor function. Indeed, CAR T cells manufactured with dex conferred a survival advantage to tumor bearing mice, which was increased in the presence of exogenous huIL7 (**Figure 6, Figure S5**). We and other have previously shown that activation using CD3 during *ex vivo* culture of CAR T cells downregulates IL7Rα (**Figure 2**) (33-35). While previous efforts have been made to upregulate IL7Rα on CAR T cells through i.e. constitutive expression of IL7Rα, there is concern has these methods have the potential to be tumorigenic (12, 13, 36-38). Because dex-mediated IL7Rα expression on CAR T cells is transient (**Figure 4; Figure S3**), possibly due to the 36-72-hour half-life of dex (39), we propose that the addition of dex to the CAR T cell manufacturing platform is a safe and clinically translatable method to increase IL7Rα. By generating IL7Rα+ CAR T cells with dex, we can thus leverage the increase in IL-7 induced by lymphodepleting chemotherapy that is standardly given 2-3 days prior to CAR T cell infusion (10, 40-44). Moreover, the benefit of dex during CAR T cell manufacturing extends beyond IL7Rα, as we observed many different pathways modulated by dex reverted back to similar trends as pre-treatment levels by day 23-24 (**Figure 4**).

Dex is used to treat several different kinds of cancer, such as multiple myeloma and glioblastoma (45, 46), as well as to treat immune-related toxicities associated with CAR T cell therapy. While there is concern that dex may decrease the function and persistence of CAR T cells (18, 19), we found that dex enhanced the efficacy of CAR T cells in our model (**Figure 7, Figure S7**). These results are supported by the recent clinical observation that CAR T cells had better expansion and persistence in patients who received dex compared patients who did not receive dex (20). These findings have the additional implication that dex may enhance CAR T cell function for other tumors, including solid tumors.

Collectively, we provide evidence that dex exerts a positive effect on CAR T cells that yields CAR T cells with superior antitumor function and persistence. Mechanistically, we found that dex improved CAR T cell function by upregulating genes in multiple pathways, including upregulation of IL7Rα. We propose that this driving mechanism can be leveraged in the CAR T cell manufacturing process to counteract the activation-induced downregulation of IL7Rα during *ex vivo* expansion. Moreover, when combined with IL-7 *in vivo*, dex enhanced CAR T cell persistence and anti-tumor activity. Most importantly, these effects were reversible and, therefore, are unlikely to induce uncontrolled CAR T cell expansion. In summary, our data supports further exploration for the use of glucocorticoids during immunotherapy and provides rationale to interrogate whether it is necessary to taper off ongoing steroid treatment prior to CAR T cell infusion.

## Methods

### Cell Lines

Human lymphoblastoid cells (LCL) were generated as previously described by transforming PBMCs with Epstein Barr virus (47). OKT3-expressing LCLs (LCL-OKT3) were generated by transfecting OKT3-2A-Hygromycin_pEK plasmid as previously described and selected for with 0.4mg/mL hygromycin (InvivoGen, ant-hg-1) (12). LCL and LCL-OKT3 cells were maintained in RPMI 1640 and 10% heat-inactivated FBS (Hyclone, SH30070.03HI). KG-1a cells were cultured in IMDM (Life Technologies, 12440-053) and 10% heat-inactivated FBS. MM.1s cells were cultured in RPMI 1640 with 10% FCS. To generate firefly luciferase+GFP+MM.1S (fflucGFPMM.1S), MM.1S cells purchased from ATCC were transduced with lentiviral vector encoding eGFP-ffluc. SUP-B15 (ATCC, CRL-1929) cells were maintained in IMDM and 20% FBS. To generate firefly luciferase(ffluc)+GFP+ cell lines, SUP-B15 cells were transduced with an eGFP-fflucepHIV viral vector and sorted for 100% purity. Chinese Hamster Ovary (CHO) (ATCC) cells were transduced with hIL7_pIRESpuro3 plasmid to produce human recombinant IL-7. CHO-IL7 cells were maintained in 50/50 DMEM/Ham’s F-12 (Corning, 10092CVR) with 10% heat-inactivated FBS and 10ug/ml puromycin (InvivoGen, anti-pr-1)

### T Cell Isolation

Healthy donor blood was obtained from City of Hope Blood Donor Center under protocols approved by City of Hope Institutional Review Board (IRB). To isolate PBMCs, blood was resuspended in PBS/2%FBS/EDTA and separated using Ficoll-Paque Plus (Cytiva, 17144002) density gradient centrifugation in SepMate50 tubes (StemCell Technologies, 85450), followed by two washes in PBS/2%FBS/EDTA. To isolate naïve and memory T cells (Tn/mem), PBMCs were resuspended in autoMACS Running Buffer (Miltenyi Biotech, 130-091-221) and up to 5 × 10^9^ cells were incubated with anti-CD14 microbeads (Miltenyi Biotech, 130-050-201) to eliminate CD14+ monocytes, and anti-CD25 microbeads (Miltenyi Biotech, 200-070-211) to eliminate CD25+ regulatory T cells, for 30 minutes on ice. CD14+CD25+ cells were immediately depleted using the DEPLETES program on autoMACS Pro Separator (Miltenyi Biotech, 130-090-273), according to the manufacturer’s protocol. The negative unlabeled cells were resuspended in autoMACS Running Buffer and anti-CD62L microbeads (Miltenyi Biotech, 170-076-700), incubated for 20 minutes on ice, and immediately subjected to the POSSELD enrichment program on autoMACS, according to the manufacturer’s protocol. For Tcm isolation, PBMCs were incubated with anti-CD14, anti-CD25, and anti-CD45RA (Miltenyi Biotech, 130-045-901) microbeads and depleted using the DEPLETES program on autoMACS. The unlabeled negative fraction was labeled with anti-CD62L microbeads and enriched with POSSELD program on autoMACS.

To generate EBV-specific T cells, purified Tcm (CD45RO+CD62L+) and Tem (CD45RO+CD62L-) cells were stimulated with 8000 rad-irradiated autologous LCL cells at 4:1 (responder/stimulator) ratio weekly for three weeks. Resultant EBV-specific T cells were then further expanded with a rapid expand method as previously described (23).

### Generation of CAR T cells

Isolated PBMC, Tn/mem, or Tcm cells were stimulated with GMP Human T-expander CD3/CD28 Dynabeads™ (Dynal Biotech Cat#11141D) at a ratio of 1:3 (T cell:bead). The following day, T cells were transduced with CD19R(EQ):CD28:ζ/EGFRt or CS1R(HL-CH3):41BB:ζ/EGFRt constructs (12, 48) at MOI=1 in RPMI 1640 containing 10%FBS and 5μg/ml protamine sulfate (Fresenius Kabi, 22905), 50U/mL rhIL-2 (Novartis Pharmaceuticals, NDC0078-0495-61), and 0.5 ng/mL rhIL-15 (CellGenix, 1013-050). Seven days after stimulation, CD3/CD28 Dynabeads™ beads were removed using a DynaMag™-5 Magnet (Invitrogen, 12303D) and plated to a concentration of 7×10^5^ cells/mL in RPMI 1640/10% FBS and 50U/mL rhIL-2 and 0.5 ng/mL rhIL-15 to rest for 48 hours. CAR T cells were cultured for 16-23 days before being used for *in vitro* experiments or being frozen in CryoStor (Biolife Solutions, 205102) for mouse experiments. For dex titration studies, 0.1μM, 1μM, or 10μM dex (Sigma, D4902) was added to the cell culture media at Day 9 only or Day 9, 12, and 14. For extended growth, cells were washed on Day 16 and grown without residual dex. IL-2 and IL-15 were supplemented every other day. Where indicated, IL-7 (R&D Systems, 207-IL-025) was supplemented to the media on Days 9, 12, and 14.

### Antibodies and Flow Cytometry

For surface staining, cells were incubated with fluorochrome-conjugated monoclonal antibodies (mAbs) to CD3 (BD Bioscience, 563109, 557832), CD4 (BD Bioscience, 557852, 340133), CD8 (BD Biosciences, 348793), CD62L (BD Biosciences, 341012), CD127 (Biolegend, 351319), EGFR (Biolegend, 352906), LAG3 (LSBio, LS-B2237), PD1 (Invitrogen, 47-2799-42), TIM3 (R&D Systems, FAB2365P), CD45RA (BD Biosciences, 555488), CD45RO (BD Biosciences, 561137), CD45 (BD Biosciences, 340665), CD107a (BD Biosciences, 555800). Cells were resuspended in FACS buffer, which consists of HBSS (Gibco, 14175095), 2% FBS and NaN3 (Sigma, S8032), and incubated with antibodies at 4°C in the dark. Following a washing step with FACS buffer, DAPI (Invitrogen, D21490) was added for viability staining before analysis.

For intracellular staining, cells were stained with FVD Viability Viogreen (Thermo Fisher Scientific, 65-0866-18) at 1:1000 dilution for 15 minutes at 4°C. The cells were washed with FACS buffer, fixed and permeabilized with Cytofix/Cytoperm Plus (BD Bioscience, 555028) for 20 minutes at 4°C, then stained with intracellular antibody for IFN-γ (BD Biosciences, 557643) at 4°C for 20 minutes. Flow cytometry was performed using MASCQuant Analyzer 10 (Miltenyi Biotech, 130-096-343), according to the manufacturer’s protocol. Flow cytometry results were analyzed using FCS Express 7 Research Edition.

### Degranulation

CAR T cells (2×10^5^) were cocultured with LCL (2×10^5^) cells, BD GolgiStop^™^ protein transport inhibitor (BD Bioscience, 554724), and antibody for CD107a (BD Bioscience, 555800) in the dark for 6 hours at 37°C. OKT3 and KG-1a cells were used as positive and negative controls, respectively. CD107a expression was determined by flow cytometry.

### NanoString Gene Expression Analysis

RNA preparation was performed according to protocol for the nCounter FLEX™ system. Raw data was processed with nCounter Advanced Analysis software (version 2.0.134) for pathway score analysis following manufacturer’s instructions. Pathways with similar changes (either upregulation or downregulation) in PBMC-derived CD19-CAR T cells and Tn/mem-derived CD19-CAR T cells at indicated timepoints were included. For gene expression analysis, raw data was first processed using nSolver 4.0 Analysis software (NanoString). Gene expression counts were normalized to positive controls and selected housekeeping genes (counts > 100 and %CV > 40 per manufacturer’s suggestions). After normalization, gene expression between cells treated with dex (Dex+) versus without dex (Dex-) at specific timepoints were compared. The formula (Count_Dex+_ - Count_Dex-_)/Count_Dex-_ was applied. Genes were then grouped into upregulated group (> 0), no change group (= 0), and downregulated group (< 0) based on the formula results. Genes in different groups within PBMC-derived CAR T cells and Tn/mem-derived CAR T cells were excluded. To further exclude genes with minor changes, the %CV cutoff value of 20 was applied based on the highest %CV from the selected housekeeping genes after normalization. The selections were done using R 3.6.1 and Excel (Microsoft).

### *In vivo* xenograft model

All animal experiments were performed under protocols approved by City of Hope Institutional Animal Care and Use Committee (IACUC). For all studies, 5×10^5^ fflucGFP SUP-B15 cells or 2×10^6^ MM.1S cells were intravenously (i.v.) or intraileally (i.t.) injected into each 6-8-week old female NSG mouse on Day −5. For dex treatment on CD19-CAR or CS1-CAR T cells, 1 × 10^6^ CAR T cells (PBMC or Tn/mem), with or without *ex vivo* dex treatment, were injected i.v. into mice. For *in vivo* dex treatment, 1 × 10^6^ Tcm-derived CD19-CAR T cells were injected i.v. and 10mg/kg Dex was injected intraperitoneally (i.p.) every 48 hours. For IL-7 injections, IL-7-producing CHO cells were irradiated at 8000 rads and injected i.p. every 48 hours. Tumor burden was monitored by live mice imaging using LagoX optical imaging system (Spectral Instruments Imaging). For imaging, mice were injected i.p. with XenoLight D-luciferin potassium salt (Perkin Elmer, 122799). Images were analyzed using Aura Imaging Software (Spectral Instruments Imaging). Humane endpoints were used to determine survival. Upon euthanasia bone marrow, blood, and spleen were harvested for flow cytometry analysis.

### Statistics

Analysis was performed using Prism (GraphPad Software Inc.). The non-parametric Mann-Whitney test was applied to group comparisons and Friedman test was applied to comparison of growth curves. Kaplan-Meier method were applied to survival function estimation and group comparisons. P-values of ≤0.05 were considered statistically significant.

### Study Approval

NOD-scid IL2Rgammanull (NSG) were purchased from The Jackson Laboratory. The NSG breeding colony was maintained by the Animal Resource Center at the City of Hope. Mice were housed in a pathogen-free animal facility according to institutional guidelines. All animal studies were approved by the Institutional Animal Care and Use Committee (IACUC: 21034). Healthy donor blood was obtained from City of Hope Blood Donor Center under protocols approved by City of Hope Institutional Review Board (IRB 09025).

## Acknowledgement

We thank Dr. Holly Yin at City of Hope for assisting Nanostring analysis. The Small Animal Imaging Core is supported by the National Cancer Institute of the National Institutes of Health (P30CA033572); Internal Institute funds.

## Author Contributions

X.W. and S.J.F. designed and directed the study, funding acquisition. A.M. and R.U. conducted, analyzed, organized the data, wrote manuscript, and contributing equally to this manuscript. E.T, K.J., D.A, and V.V. performed experimental work. S.H.L. and C.H. conducted NanoString data analysis. M.C.C. reviewed and edited the manuscript. S.J.P. provided design of solid tumor experiments and reviewed the manuscript. All authors read and approved the final version.

