## Supplemental Figures for "Dexamethasone Enhances CAR T Cell Persistence and Function by Upregulating Interleukin 7 Receptor"

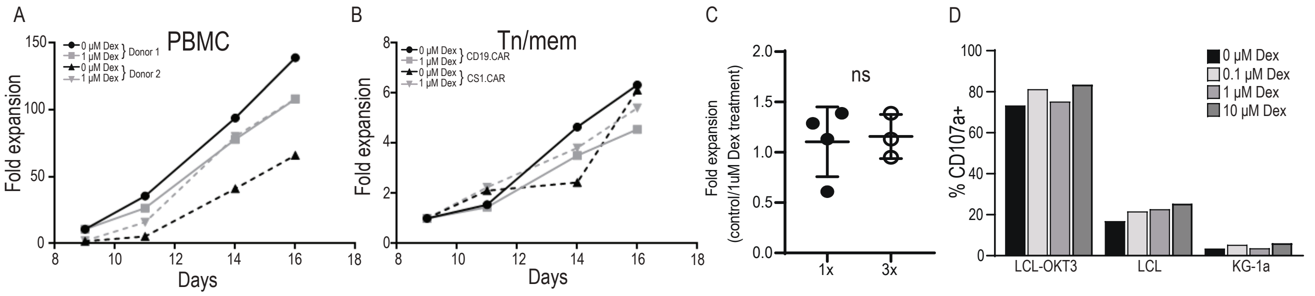


**Supplemental Figure 1. Growth and function of CAR T cells treated with various concentrations of dex.** Proliferation of PBMC (n=2) **(A)** or Tn/mem (n=2) **(B)** CAR T cells grown in the presence or absence of a single 1μM dex treatment. **(C)** Cumulative data of fold expansion from all donors treated with either single or three treatments of 1μM dex. Data was normalized to untreated CAR T cells. **(D)** CD107a degranulation by Tn/mem-derived CAR T cells pre-treated with various dex concentrations (0.1, 1, 10μM).

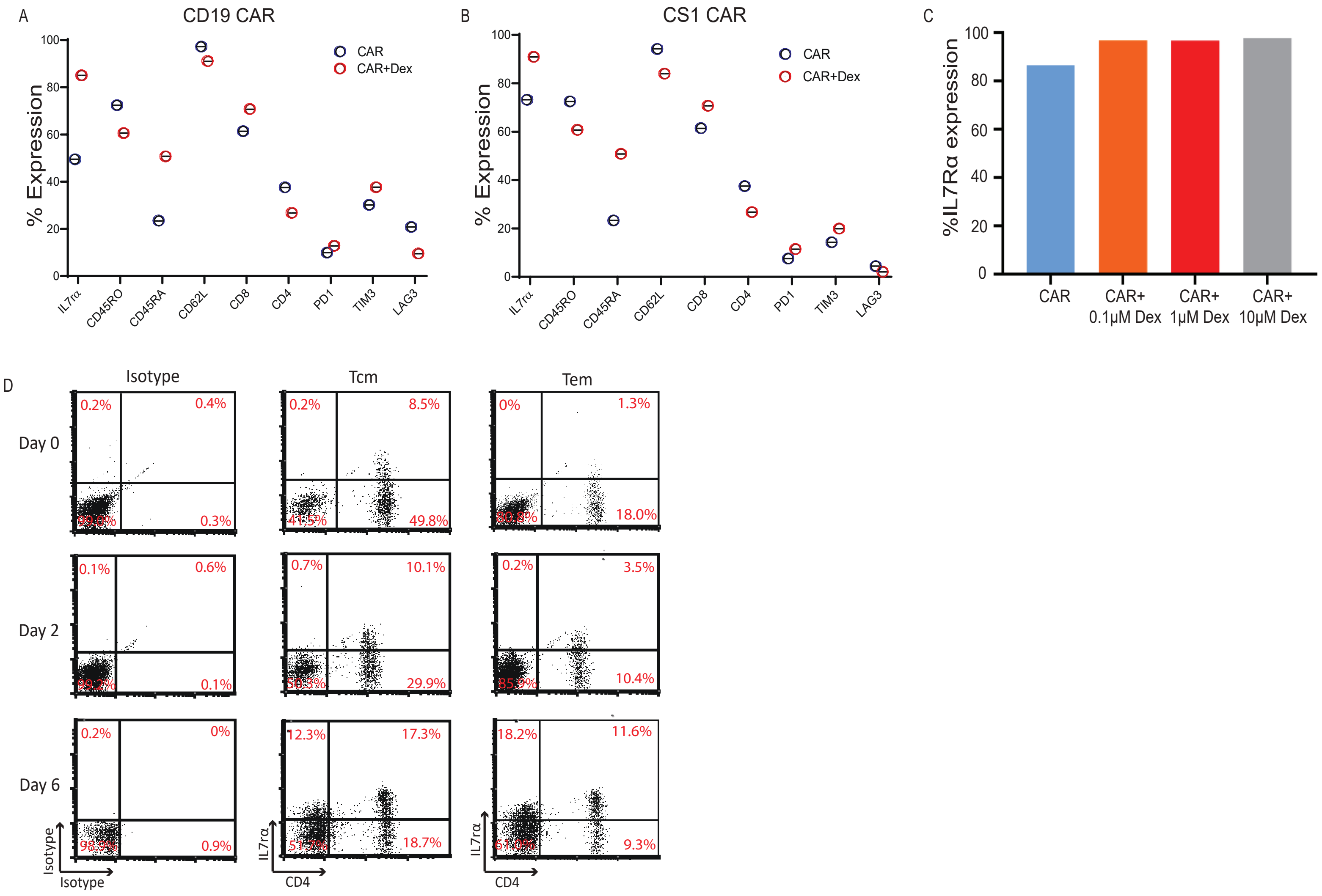


**Supplemental Figure 2. Expression of IL7Rα is consistent in dex-treated CAR T cells regardless of CAR constructs, T cell populations, or dex concentrations.** Phenotype of Tn/mem-derived anti-CD19 **(A)** or CS1 **(B)** CAR T cells in the presence or absence of 1μM dex treatment. Expression of IL7Rα at day 16 on Tn/mem CAR T cells in the presence or absence of various concentrations of dex (0.1, 1, 10μM) **(C)**. REM expanded EBV-specific Tcm and Tem cells were treated with 1uM Dex and levels of IL7Rα were analyzed at different time points after Dex treatment **(D)**.

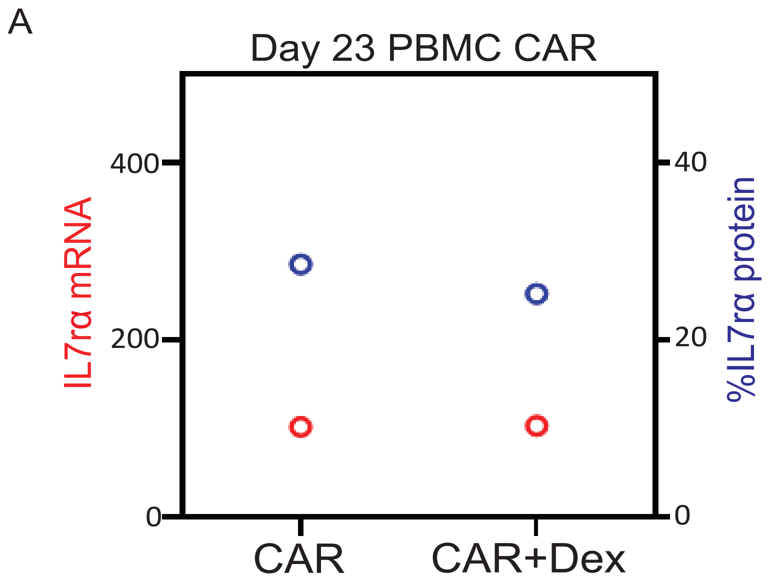


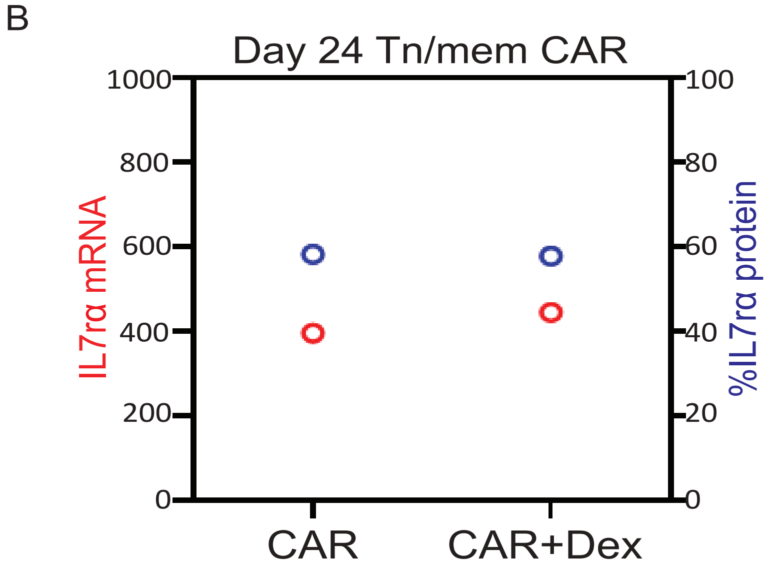


**Supplemental Figure 3. Reversible patterns of dex-upregulated IL7Rα in CAR T cells.** CD19-CAR T cells were treated with 1μM dex (1x) on day 9, dex was removed from culture on day 16, and cultures were extended to day 23 (PBMC CAR) **(A)** and 24 (Tn/mem CAR) **(B)**. Protein IL7Rα expression was analyzed with flow cytometry and mRNA of IL7Rα was analyzed with Nanostring technology.

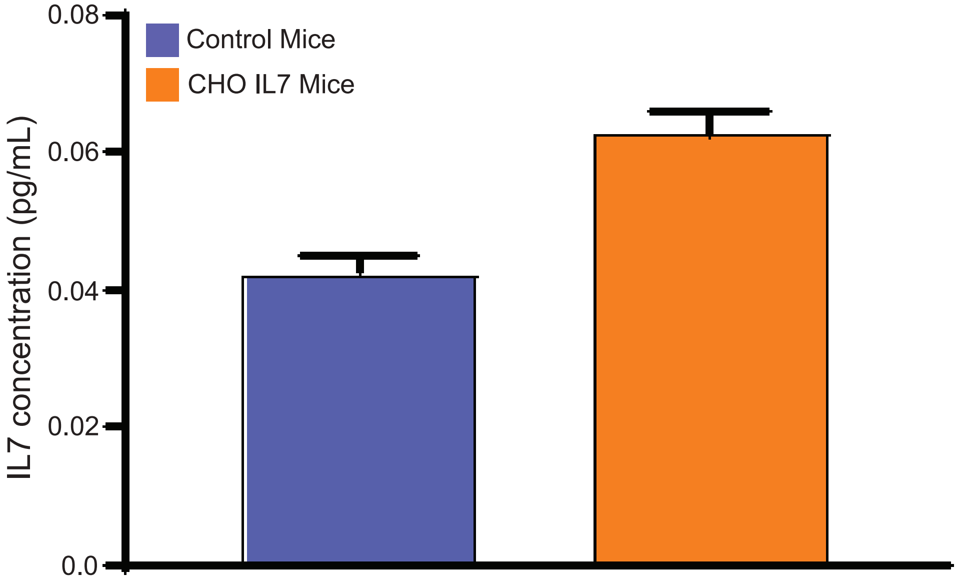


**Supplemental Figure 4. Serum concentration of human IL7 in mice injected with IL7-secreting CHO cells.** Serum from mice treated with IL7-producing CHO cells was collected 24 hours post-IL7-CHO cell injection. Serum from untreated mice was used as a control. Human IL-7 was measured with ELISA.

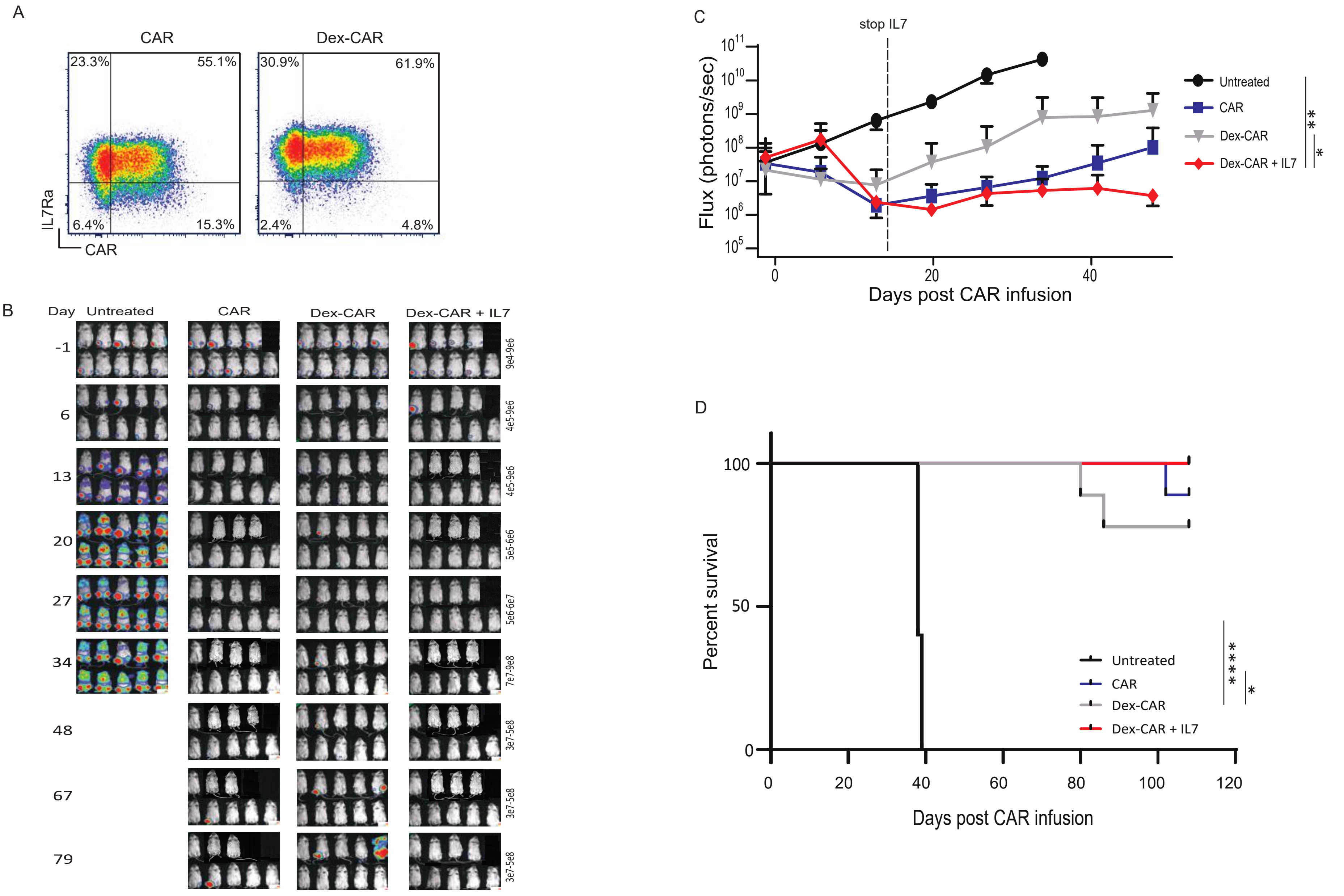


**Supplemental Figure 5. Dex-treated anti-CS1 CAR T cells in MM.1s xenograft model.** $2 \times 10^6$ multiple myeloma (MM.1s) cells expressing GFP and firefly luciferase (GFPffluc+) were intratibially (i.t.) injected into NSG mice. Five days later, when tumor engraftment was confirmed, $1 \times 10^6$ CAR T cells with or without dex ex vivo treatment were injected i.v. Mice were injected intraperitoneally (i.p.) with 8000 rads irradiated human IL-7-producing CHO cells ($10 \times 10^6$) every 48 hours for six injections. IL7Rα expression on input cells **(A)** after 16 days expansion (7 days in dex). Biophotonic tumor imaging for tumor signal and tumor burden was measured in Flux (photons/sec) by bioluminescent imaging and was evaluated weekly **(B, C)**. Kaplan-Meier survival curve **(D)**.

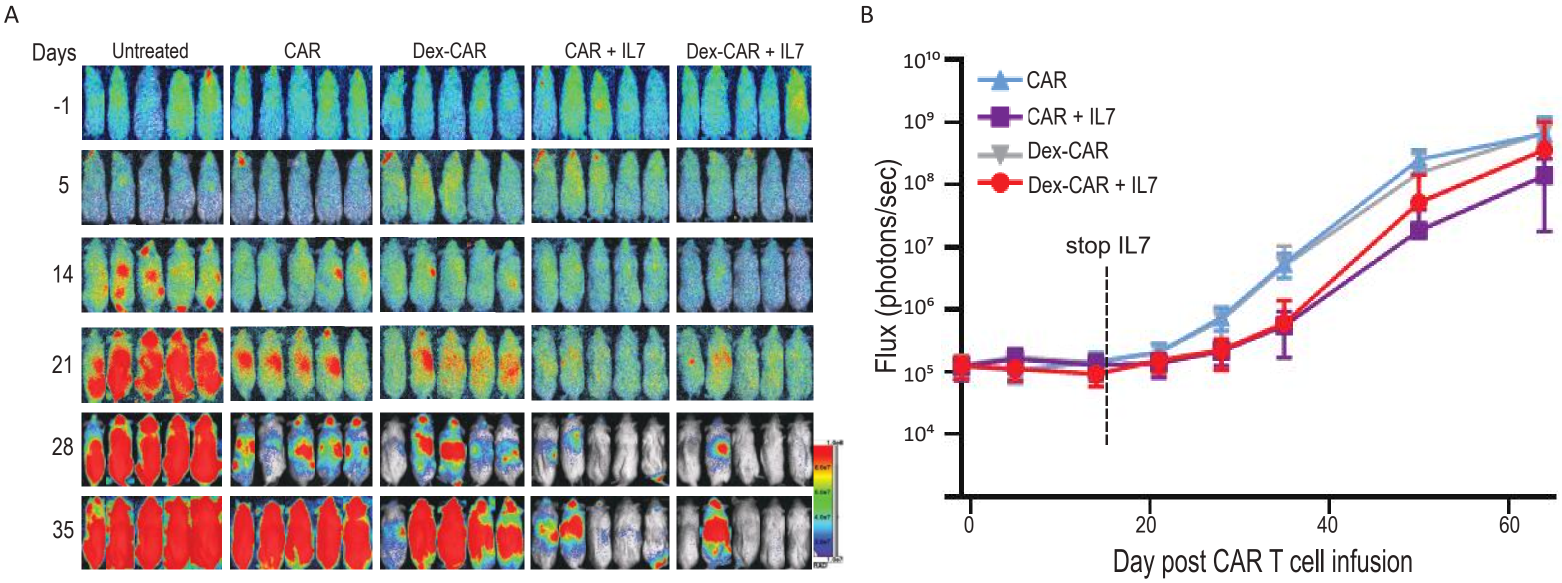


**Supplemental Figure 6. Day 23 expanded Dex-treated anti-CD19-CAR T cells in SupB15 xenograft model.** 0.5 x $10^6$ acute lymphoblastic leukemia cell line (SupB15) cells expressing GFP and firefly luciferase (GFPfflucȧ+) were inoculated into NSG mice intravenously (i.v.) on Day -5. After confirmation of engraftment, 1 x $10^6$ day 23 (7 days in dex, 7 days removed from dex) expanded CAR T cells were adoptively transferred into tumor-bearing mice i.v. Mice were injected intraperitoneally (i.p.) with 8000 rads irradiated human IL-7 producing CHO cells (10 x $10^6$) every 48 hours for six injections. Biophotonic imaging was used for tumor signal monitoring **(A)** and tumor burden was measured in Flux (photons/sec) by bioluminescent imaging and was evaluated weekly **(B)**.

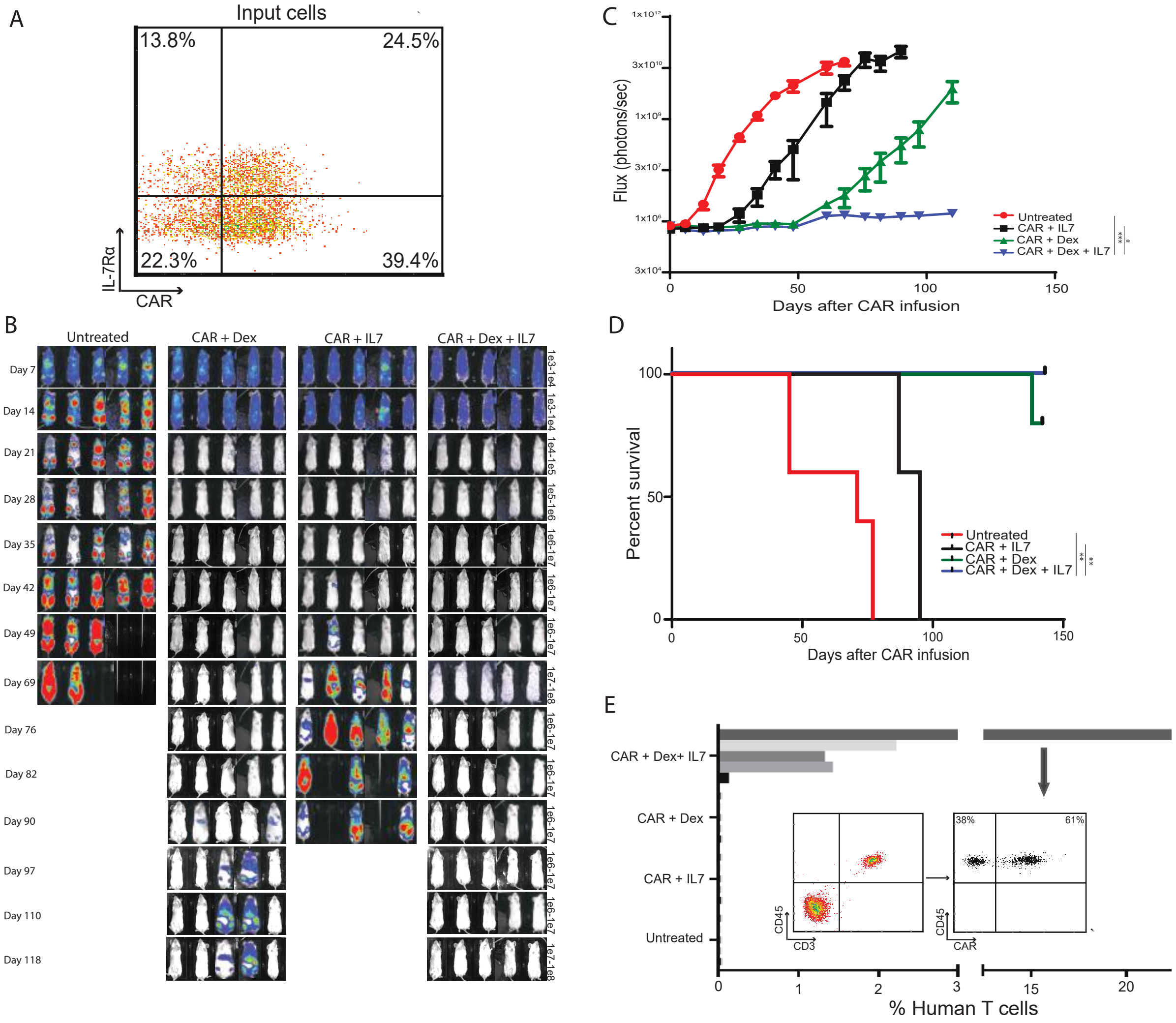


**Supplemental Figure 7. Combination of CD19-CAR T cells with dex and IL7 against SupB15 xenograft model.** Mice (n=5) were engrafted with 0.5 x $10^6$ SupB15GFPffluc+ cells i.v. on Day -5. 1 x $10^6$ CD19-CAR T cells were injected i.v. Dex was given 10mg/kg s.c. every other day for 4 weeks and then once a week afterwards. 10 x $10^6$ 8000 rads-irradiated human IL7-secreting CHO cells were injected i.p. every other day for 4 weeks and then once a week afterwards. CAR expression on input cells **(A)**. Biophotonic tumor imaging for tumor signal and tumor burden was measured in Flux (photons/sec) by bioluminescent imaging and was evaluated weekly **(B, C)**. Kaplan-Meier survival curve **(D)**. On Day 21, blood was collected through retroorbital bleeding and analyzed for CAR expression by flow cytometry **(E)**. **P<0.01, ***P<0.001.
